# Combining 16S rRNA gene variable regions enables high-resolution microbial community profiling

**DOI:** 10.1101/146738

**Authors:** Garold Fuks, Michael Elgart, Amnon Amir, Amit Zeisel, Peter J. Turnbaugh, Yoav Soen, Noam Shental

## Abstract

**Background:** Most of our knowledge about the remarkable microbial diversity on Earth comes from sequencing the 16S rRNA gene. The use of next-generation sequencing methods has increased sample number and sequencing depth, but the read length of the most widely used sequencing platforms today is quite short, requiring the researcher to choose a subset of the gene to sequence (typically 16-33% of the total length). Thus, many bacteria may share the same amplified region and the resolution of profiling is inherently limited. Platforms that offer ultra long read lengths, whole genome shotgun sequencing approaches, and computational frameworks formerly suggested by us and by others, all allow different ways to circumvent this problem yet suffer various shortcomings. There is need for a simple and low cost 16S rRNA gene based profiling approach that harnesses the short read length to provide a much larger coverage of the gene to allow for high resolution, even in harsh conditions of low bacterial biomass and fragmented DNA.

**Results:** This manuscript suggests Short MUltiple Regions Framework (SMURF), a method to combine sequencing results from different PCR-amplified regions to provide one coherent profiling. The *de facto* amplicon length is the total length of all amplified regions, thus providing much higher resolution compared to current techniques. Computationally, the method solves a convex optimization problem that allows extremely fast reconstruction and requires only moderate memory. We demonstrate the increase in resolution by *in silico* simulations and by profiling two mock mixtures and real-world biological samples. Reanalyzing a mock mixture from the Human Microbiome Project achieved about two-fold improvement in resolution when combing two independent regions. Using a custom set of six primer pairs spanning about 1200bp (80%) of the 16S rRNA gene we were able to achieve ~100 fold improvement in resolution compared to a single region, over a mock mixture of common human gut bacterial isolates. Finally, profiling of a *Drosophila melanogaster* microbiome using the set of six primer pairs provided a ~100 fold increase in resolution, and thus enabling efficient downstream analysis.

**Conclusions:** SMURF enables identification of near full-length 16S rRNA gene sequences in microbial communities, having resolution superior compared to current techniques. It may be applied to standard sample preparation protocols with very little modifications. SMURF also paves the way to high-resolution profiling of low-biomass and fragmented DNA, e.g., in the case of Formalin-fixed and Paraffin-embedded samples, fossil-derived DNA or DNA exposed to other degrading conditions. The approach is not restricted to combining amplicons of the 16S rRNA gene and may be applied to any set of amplicons, e.g., in Multilocus Sequence Typing (MLST).

## Background

Bacteria are the most diverse domain on our planet [1], playing a crucial role in numerous ecosystems as well as an inseparable role in human health and disease. With the advent of next generation sequencing technologies, molecular identification has replaced traditional culturing, allowing a window into the diversity of microbial communities. Most of these studies are based on sequencing of the 16S rRNA gene, which has several highly conserved regions interleaved with variable regions. Conserved regions serve as 'anchors' for designing polymerase chain reaction (PCR) primers while sequencing the variable regions identifies the bacteria.

Such PCR primers are required to be 'universal', *i.e.,* amplify most bacterial 16S rRNA gene sequences, while maximizing the phylogenetic resolution, *namely,* the ability to distinguish among different bacteria based on the amplified sequence (for a comparison between different regions see [2] and references therein). However, the main design criterion for many current 16S rRNA gene primers is to increase the universality under the constraint of amplifying a region that matches the read length available by the sequencing platform *(e.g.* V3-V4 primers for Illumina paired-end sequencing [3–6]). Hence resolution is limited since many bacteria may share the same short amplified sequence.

High resolution identification of bacteria in mixtures may prove highly useful in subsequent analysis, especially when follow-up studies require isolation and monoculture experiments. The basic aim is, therefore, to allow an exact identification of each bacterium in a mixture, or at least minimize the *ambiguity, i.e.,* the number of bacteria that share the same 16S rRNA gene 'footprint'.

## Using short reads over a long amplicon to increase resolution

The main reason for matching the amplicon length to the read length is technical and not biological. In recent years several computational methods that allow higher resolution profiling based on an amplicon longer than the read length have been proposed [7–12]. These methods receive short reads that originate from one long amplified region, and solve some optimization problem that seeks a combination of bacteria that best explains the set of reads in the experiment. Different methods vary in their fine details, representation, applied cost function and optimization method. This approach, which we term the “single long-region framework”, suffers from several drawbacks:

i. *Non-standard 16S rRNA gene sample preparation:* Prior to sequencing, a long amplicon requires shearing *(e.g.,* by sonication) followed by size selection and adaptors' ligation. Apart from being quite labor intensive and time-consuming, both size selection and ligation steps cause significant material loss that could be detrimental when detecting low frequency bacteria in complicated biological mixtures. An additional PCR step is required to compensate for such material loss, hence additional PCR bias introduced.
ii. *Primers are far from being universal:* Current primers for amplifying the whole 16S rRNA gene (e.g., 8F and 1492R [13]) significantly improve resolution, compared to sequencing a short region, however they are highly non universal. To increase universality some mix of primer pairs is required.
iii. *Sequencing fragmented DNA* may not be feasible: Increasing resolution via sequencing a long amplicon implicitly assumes that available DNA molecules are long enough. Although this is often the case, there are conditions in which molecules are fragmented, *e.g.,* when DNA is extracted from Formalin-fixed and Paraffin-embedded (FFPE) blocks, fossil-derived DNA, or DNA exposed to harsh environments (radiation, reducing agents, etc.). An average length of a DNA molecule in this case may be shorter than needed for amplifying a long region, and thus amplification may not be possible. Moreover, sometimes even standard primers may not be suitable since the molecules' lengths are too short (if the average molecule length is about 200bp then also custom V3-V4 primers may not be applied).

## Short multiple-regions framework

Here we propose a different approach that allows high resolution profiling. The method is based on independent PCR amplification and sequencing of several (actually any number) of regions along the 16S rRNA gene, which are computationally combined to provide a joint estimate of the microbial community composition (Figure 1). We term this method “Short MUltiple Regions Framework” (SMURF). Since several regions are amplified, the *de facto* amplicon is large allowing for high resolution similar to the single long-region framework, while having the following advantages:

i. *Availability of custom primers and standard sample preparation:* SMURF can be applied to any combination of common primer pairs (e.g. V1-V3, V3-V5, V4 etc.) without the need for designing custom long-amplicon primers. Also, DNA shearing, size selection and ligation steps needed in the long-amplicon case are not required. Basically, the approach allows high resolution without the need for modifying experimental procedures (namely library preparation is independently performed for each primer pair, as in standard protocols).
ii. *'Mix-and-match' primers to increase primers' universality and resolution:* Increasing universality and resolution in SMURF amounts to amplifying additional regions until the required universality and resolution are achieved. Rather than aiming to improve universality [14] or searching for *the* single and optimal set of primers [2], SMURF allows to combine any set of 'sub-optimal' primers to achieve both superior universality and resolution compared to each of these primers applied by itself. Results would evidently depend on the properties of the chosen set of primers, and therefore a wise selection would yield more accurate profiling.
iii. *SMURF allows sequencing fragmented DNA:* The 'building blocks', i.e., the amplicons, are short and thus allow amplification also in case of highly fragmented DNA.
iv. *Multiple regions tend to 'average' PCR bias:* PCR amplification may have a different efficiency for different bacteria [15, 16], which results in considerable differences in profiling depending on primer choice [17–20]. Moreover, Gohl et al. [21] have shown that the *same* primer pair may yield different profiling results depending on the specific library preparation protocol. Since SMURF reconstruction is performed based on several amplified regions, and assuming that biases are independent in every region, reconstruction tends to average the effects of such biases.
v. *Algorithmic advantages:* The mathematical formulation of SMURF is analogous to the one formerly suggested for the long-region methods [7, 12]. However, there are several computational advantages of SMURF that stem from combining short regions. First, long-region methods require aligning reads to reference genomes before estimating bacterial frequencies. This time-consuming and error-prone step (e.g. via Bowtie [22] or BWA [23]) is not needed in SMURF since each read is readily associated with its primer and then matched to the corresponding region's k-mers database (a k-mer in this context is a bacterial sequence from the amplified region whose length matches the read length). Second, data representation in this case is much more efficient. The number of k-mers representing a bacterium is orders of magnitude smaller than for the long region framework (a bacterium is represented by one or two k-mers per region as opposed to the long region framework, where the number of k-mers is equal to the region's length). Consequently, optimization does not require handling huge matrices, which together with fixed point iteration formula results in an extremely fast reconstruction. The third computational advantage is that the SMURF cost function is convex and thus convergence to a global optimum is guaranteed.

**Figure 1:**
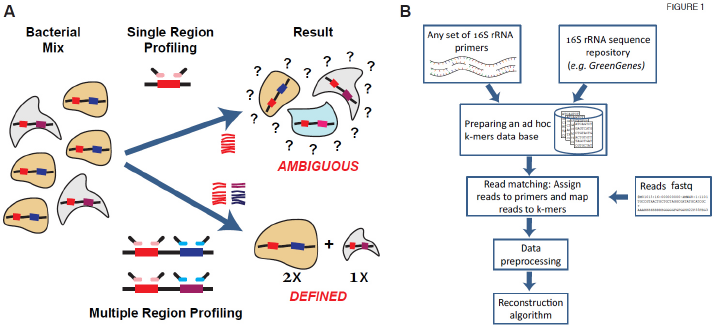
A Schematic description of SMURF. (**A**) A comparison between single and multiple region profilings. **(B)** The SMURF flow diagram describing the internal steps of a typical analysis.

The manuscript presents the improvement in resolution achieved by SMURF and demonstrates the advantages of detecting near full-length 16S rRNA gene sequences. The work is divided into several parts: (i) First, using *in silico* simulations we show a marked increase in resolution as a function of the number of amplified regions; (ii) Second, the correctness of the method is assessed and verified via analysis of several mock mixtures. In the first experiment, a mock mixture was sequenced using a new set of six primer pairs each amplifying about 200bp of the 16S rRNA gene. The experiment allowed us to confirm the increase in resolution as a function of the number of applied primer pairs. Next we re-analyzed a mock mixture from the Human Microbiome Project combining reads originating from two regions (V1-V3 and V3-V6). Apart from correct reconstruction and improvement in resolution when combining two regions, this experiment displayed the applicability of SMURF to available data acquired via standard protocols; (iii) Finally a real-world experiment characterizing a dynamic change occurring in the complex *D. melanogaster* microbiome was analyzed. We demonstrate the correctness of SMURF's results and describe how high resolution profiling was essential for efficient downstream analyses.

## Methods

Steps in SMURF are detailed (Figure 1B), followed by a description of simulations and experiments performed (a Matlab code of SMURF is available at https://github.com/NoamShental/SMURF).

## Database preparation

*Database:* The basic database used was the Greengenes (GG) 16S rRNA gene database (May 2013 version) [24] (other databases, *e.g.*, SILVA [25], can also be used as described in the SMURF software package). Sequences containing *ambiguity* were duplicated by all possible sequence values (up to 64 combinations per sequence; sequences that contained more than three ambiguous nucleotides were discarded). Consequently, the basic database consisted of 1,402,801 unique 16S rRNA gene sequences. *Preparing an ad hoc, protocol specific, database:* Given the sequencer's read length, *k* and each region's primer pair, a list of all possible k-mers in each region was generated based on the basic database. A bacterium was termed 'amplified' in a specific region if each of the forward and reverse primers had at most two mismatches with the bacterial sequence. Data was presented using a database matrix, *M*, where *M*_*hj*_ is the probability of observing the k-mer *h* in bacterium *j*, *M_hj_* = *Pr*(*kmer* = *h\bacterium* = *j*). The matrix *M* holds the information of all regions (*i.e.*, for all / bacteria and = possible k-mers). In (the rare) case that the same k-mer appears in two regions, *M* holds a separate line for each.

For paired-end sequencing each bacterial sequence may contribute a single k-mer per region (the k-mer is a concatenation of two bacterial sequences of length *k* that begin from the two ends of the amplicon). Hence, *M_hj_* = 1/*R_j_*, where *R_j_* is the number of regions amplified for bacterium *j*, which also equals the number of k-mers originating from bacterium *j*. For singleend sequencing there are two possible k-mers per amplified region (one from each amplicon ends), *i.e.*, *M_hj_* = 0.5/*R_j_*. Hence the maximal matrix size is the number of bacterial sequences in the database multiplied by twice the number of regions, which is orders of magnitude smaller than for single long-region framework, which greatly simplifies algorithm complexity and run time (*e.g.*, the analogous matrix *M* in COMPASS [9] contains ~100 fold more rows).

## Read matching

Reads were assigned to a region by the primer sequence, and hence 'alignment' amounts to counting the number of mismatches between a read and the list of k-mers in that region (compared to applying complicated and costly alignment algorithms such as Bowtie required in the long region approach). A read is termed 'matched' to a k-mer if there were at most two mismatches between the read and the k-mer after excluding the primer sequence. Following alignment of read *i* it was assigned a probability that it originated from a k-mer *h*, E_*ih*_. Assuming a constant error probability per nucleotide *p_e_*, and independence between different locations along the read then,

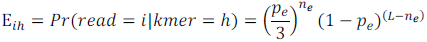

where *n_e_* is the number of mismatches between the k-mer and the read (results were almost independent of the selected value of *p_e_* (Figure S4), and thus *p_e_* = 0.005 was used [26]).

## Data preprocessing

Two filters were applied:

*Filtering low frequency reads:* Reads that appeared less than 10^-4^ times the number of reads in a region were discarded. The filter sets the lowest detectable frequency (0.01%) while significantly decreasing false positive detections resulting from reads errors. The threshold value, 10^-4^, may be lowered if the number of reads per region is high enough and bacteria whose frequencies are lower than 0.01% are sought. To decrease false positive detections in such case a read de-noising algorithm [27, 28] may be applied prior to SMURF.

*Filtering candidate bacteria:* In case no reads were aligned to a bacterium in a region for which there was a perfect match between the bacterial sequence and the primers, this bacterium was not considered in subsequent analysis.

## Reconstruction algorithm

For reconstruction we followed the maximum likelihood probabilistic framework suggested in [7, 12].

Given *M_hj_* and *E_ih_*, using the law of total probability, the probability of a read *i* given a bacterium *j*, is given by 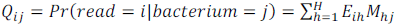. The probability of observing the read is 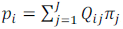, where π_*j*_ is a probability of selecting a read belonging to bacterium *j*. The likelihood of the *N* reads is given by 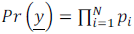, which is a convex function of the unknown read proportions vector, 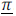 Hence, using the Expectation-Maximization algorithm [29] an iterative procedure for the estimation of 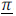 was derived.

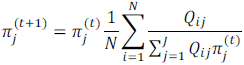

The above procedure is similar to the one used in [12], while assuming constant error probability, which results in a convex cost function and a highly efficient fixed point iteration formula.

The frequency of bacterium *j*, *x_j_*, was estimated by normalizing π_*j*_ by the number of k-mers originating from this bacterium *j*, which was simply the number of regions, *R_j_*, amplified in bacterium *j*,

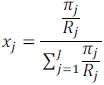

## Resolution, ambiguity and the definition of a group

SMURF results in a list of reconstructed bacteria (i.e., full-length 16S rRNA gene sequences) and their frequencies. Some of these sequences may be indistinguishable by SMURF since they share the same sequence over all amplified regions. A set of reconstructed sequences that shares the same 'footprint' over the amplified regions was defined as a *'group'.*

A bacterium may also correspond to several reconstructed *groups, e.g.,* due to several distinct 16S rRNA gene operons or because of profiling errors. Hence in order to measure resolution we define *'ambiguity'* as the effective number of 16S rRNA gene sequences that were 'assigned' to a bacterium. In case a bacterium corresponds to a single *group, ambiguity* is in fact the *group's* size. To quantify *ambiguity* we used the exponent of the Shannon entropy [30, 31]. Although *ambiguity* is actually a standard alpha diversity measure, we decided to use this term to emphasize the fact that lower values, which correspond to higher resolution, are preferred.

Resolution may not necessarily be associated with changes in taxonomy. In many cases even a single region allows species-level identification, however, sub-species may not be decided. Hence, studies that require a unique identification of full-length 16S rRNA gene sequences would benefit from applying SMURF.

## *In silico* simulations

Performance of SMURF was first evaluated by *in silico* simulations. Bacterial communities of 100 bacteria were randomly selected from the database, where their frequency followed a power law distribution (1/*x*). Profiling such a mixture is challenging since most bacteria have very low frequencies (half of the bacteria in the mixture have a cumulative frequency of 13%). The set of six primer pairs, later used in our experimental mock mixture, was applied in our simulations (the total amplicon length was about 6· 200bp). The simulated number of reads over all regions was 200,000, where the sequencing error model was adopted from [9]. Results are based on 1000 simulated communities.

*Effect of the number of regions:* Each simulated mixture was reconstructed six times, starting from a single region and adding one region at a time (the order of regions was set so as to maximize the number of unique groups in the *ad hoc* database at each step). The total number of reads was kept the same (200,000) in all cases, *i.e.,* the number of reads per region decreased when increasing the number of regions.

*Simulation performance measures:* Performance was measured by an adaptation of weighted precision and weighted recall defined in [9], quantifying the true discovery rates *(i.e.* one minus false discovery rate (1-FDR)) and true positive rate, respectively. Sequences in the simulated mixture were compared to sequences in the reconstructed mixture (comparison was performed over the whole 16S rRNA gene sequence). Whenever a perfect match occurred between a simulated bacterium *m* and reconstructed bacterium *r*, the indicator functions *I_m_* = 1 and *I_r_* = 1 were assigned. Even a single mismatch between *r* and *m* was considered as an error.

Weighted recall, *i.e.,* the probability that a bacterium belongs to the reconstructed mixture given it appeared in the simulated mixture was 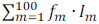, where *f_m_* was the frequency of a bacterium in the simulated mixture. Weighted precision, *i.e.*, the probability that a bacterium belongs to the simulated mixture given it appeared in the reconstructed mixture, 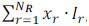, where *x_r_* is the frequency of bacterium *r* in reconstructed mixture, and *N_R_* is the total number of reconstructed bacteria.

## Experimental profiling of a mock mixture

*Primers:* To test the method a set of six primer pairs was designed (Table 1). Each primer pair was selected to maximize the number of bacteria amplified in a region, while satisfying some desired primers' characteristics *(e.g.* GC content, melting temperature, etc.). Amplicon length was about 200-250bp.

**Table 1:**
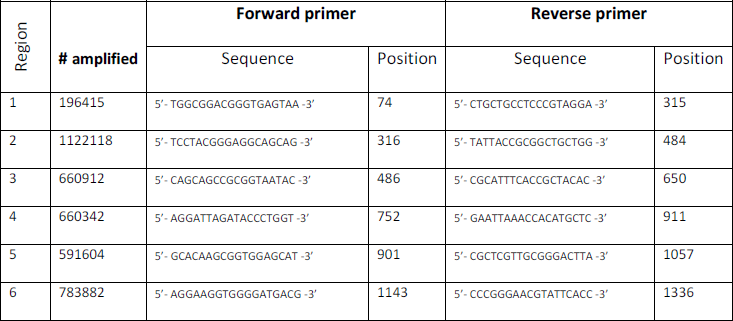
The set of six primer pairs used in this study: The number of Greengenes sequences (out of the total **1402801** sequences) that have a perfect match with each primer pair. The primers' sequences and their median start locations along the forward strand are shown. The set of primers was used for profiling our mock mixture, the *D. melanogaster* samples, and for *in silico* simulations.

*Mock mixture - Sample preparation:* Genomic DNA from ten bacterial strains: *Akkermansia muciniphila* ATCC BAA-835, *Lactobacillus casei* ATCC 334, *Bacteroides caccae* ATCC 43185, *Bacteroides fragilis* ATCC 25285, *Bacteroides ovatus* ATCC 8483T, *Eggerthella lenta* DSM2243, *Eggerthella lenta* FAA 1-3-56 (both *Eggerthella lenta* strains have the same 16S rRNA gene sequence), *Escherichia coli* BL21, *Eubacterium rectale* ATCC 33656, *and Lactobacillus plantarum* ATCC 8014, was isolated using the UltraClean^®^ Microbial DNA Isolation Kit (MO BIO Laboratories, Inc.), quantified using the Qubit™dsDNA BR Assay Kit (ThermoFisher Scientific), and mixed in equal proportions.

PCR amplification of six predefined regions was performed using the primers described above, and their products were mixed. Samples were then cleaned on column (Promega, Fitchburg, WI), concentration was measured using NanoDrop (NanoDrop Technologies, Wilmington, DE), and diluted to 50ng/ul. Subsequently, the sample went through standard Illumina library preparation, and sequenced on an Illumina HiSeq2000 sequencer using paired-end 100bp reads (the sample was sequenced together with many other samples). The number of reads per region varied between 800,000 to about 2,000,000.

## Analysis

*Preprocessing reads:* The barcodes (7bp) and the primers (18bp) were removed from the reads resulting in 76bp long reads assigned to each region. Reads were discarded in three cases (i) A Phred score of less than 30 in more than 25% of nucleotides, (ii) More than three nucleotides having a Phred score of less than 10, or (iii) When containing one or more ambiguous base calling (*e.g., 'N'*).

*Postprocessing SMURF results:* All reconstructed 16S rRNA gene sequences were classified via the Ribosomal Database Project (RDP) Sequence Match engine [32], and results were combined *(i.e.* frequencies added) for bacteria sharing the same RDP assigned species. Combined *groups* having a total frequency lower than 0.1% were discarded and the relative frequencies of the remaining *groups* was renormalized. Since both *Eggerthella lenta* strains share the same 16S rRNA gene sequence their combined frequency was shown.

*Evaluating the effect of the number of regions:* The sample was reconstructed six times, starting from a single region and adding one region at a time in the same order as described in the simulations section.

*Estimating resolution:* Resolution was separately estimated for each bacterium in the mock mixture, using the exponent of the Shannon entropy [30, 31]. For each bacterium in the mock mixture, all groups whose RDP classification matched the bacterium were considered in this *ambiguity* calculation.

## Re-analyzing Human Microbiome Project mock mixture

To further demonstrate the advantages of SMURF we examined a microbial mock community used to validate the protocols of the Human Microbiome Project (HMP) [33]. This 'even mock mixture' sample (https://www.ncbi.nlm.nih.gov/sra/SRX020130) containing 21 known bacterial strains, was independently profiled over regions V1-V3 (SRR42565 and SRR42567) and V6-V9 (SRR42570 and SRR42571). Reads from the two experimental repeats for each region were pooled together for reconstruction. The total number of reads that passed a quality filter was 56,000 and 27,000 for V1-V3 and V6-V9, respectively. SMURF was used to combine results of these two regions to display the increase in resolution. Reads were trimmed to length 170nt. Preprocessing the reads and postprocessing SMURF results were performed as described for the experimental mock mixture. To display the increase in resolution, reconstruction was performed based on each region separately and for combining the two regions.

## Profiling of a Drosophila microbiome

*Drosophila Stocks:* A *hairy-GAL4* line was obtained from the Bloomington Stock Center.

*Extracting bacterial DNA:* Flies were reared either on standard medium or medium containing 400µg/ml of the G418 toxin (Sigma). For each experiment seven to ten 3^rd^ stage larvae were collected, their gut dissected and total bacterial DNA was extracted via the “chemagic DNA Bacteria” kit (Chemagen).

*Sample preparation and sequencing:* PCR was performed using the set of six primer pairs described. Sample preparation was performed similarly to our mock mixture described above, and sequenced on the same lane. The number of reads per region varied between ~300,000 to ~1,800,000.

*Isolation and Growth of Commensal Bacteria:* Ten male flies were shaken in 1ml of PBS buffer at room temperature for 30min. 100ml of this fluid were serially diluted, spread on YPD agar plates and colonies were grown at room temperature for three days. Several colonies were picked, underwent three additional rounds of isolation, grown overnight at 30° C in liquid YPD and stocked in 35% glycerol at -80° C.

*Sanger sequencing of colonies:* The 16S rRNA gene was PCR-amplified from bacterial DNA using universal primers 8F (AGAGTTTGATCCTGGCTCAG) and 1492R (GGTTACCTTGTTACGACTT) [13]. The PCR product was cloned into PGMT plasmid, sequenced and compared to SMURF predicted sequences.

## Results

Using the set of six primers pairs, the theoretical increase in resolution as a function of the number of regions is first presented. Second, SMURF's results of both *in silico* simulations and an experimental mock mixture using this set of primer pairs are shown. Third, we display the effect of applying SMURF to re-analyze an HMP mock mixture. Finally we present the advantages of increasing resolution in detecting specific bacterial strains in real world example of D. *melanogaster* microbiome undergoing a change.

## Theoretical effect of combining short regions

To assess the potential improvement in resolution when combining several short regions, we calculated the *group* size of each bacterial sequence in the Greengenes (GG) database using our six primer pairs. Namely, the number of GG sequences that share the same sequence over the relevant regions and are thus indistinguishable were calculated (the *group's* size coincides with the *ambiguity* in this case and hence measures the resolution by which a bacterium may be identified). This procedure was repeated six times, starting from a single region and adding one region at a time as described in Methods. For comparison, data for variable region 4 (V4) is also shown (using standard forward and reverse primers: 5'-GTGCCAGCMGCCGCGGTAA-3', 5'-GGACTACHVGGGTWTCTAAT-3').

Figure 2A shows the fraction of GG sequences that belong to a *group* of up to a certain size, for one and six regions and for the V4 region (results for two to five regions were omitted for clarity and appear in Figure S1). For example, 13% of the GG sequences belong to a *group* of size one, i.e., were uniquely identifiable using a single region. Using V4 the fraction of uniquely identifiable sequences was 21%, while for six regions the fraction was doubled to about 40%, which displays the potential increase in resolution using a larger number of regions. The same phenomenon occurred also for sequences that were not uniquely identifiable and belong to larger *groups* - much more sequences belonged to *smaller groups* when using six regions compared to V4, corresponding to a significant increase in resolution. Although our single region provided inferior resolution compared to V4 (which uses a set of twenty oligonucleotides instead of just a single pair in our case), using merely two regions was sufficient for better performance than V4 (Figure S1).

## *In silico* simulations

Figure 2B shows reconstruction performance when simulating 200,000 reads from a mixture of 100 bacteria randomly selected from the GG database, having a power law frequency distribution. Weighted precision and weighted recall were plotted for one and six regions and for using V4 primers (results for two to five regions were omitted for clarity and appear in Figure S2).

Weighted recall *(i.e.* the chances that a bacterium from the simulated mixture appeared in the reconstruction) was very high in all cases. The 200,000 reads were divided among regions, and hence simulating six regions used six fold less reads per region, resulting in slightly lower performance detecting the low frequency sequences. These differences became more pronounced when simulating a lower number of reads (Figure S2). Hence, a certain minimal number of reads per region was required in order to maintain high recall performance.

Weighted precision *(i.e.* the chances that a bacterium from the reconstructed community appeared in the original mixture) greatly improved when using six regions compared to a single region and to V4. Improvement in precision was mainly achieved by a reduction in the *groups'* sizes, due to the longer effective amplicon. The probability of correctly detecting a bacterium which is uniquely identifiable, i.e., belongs to a *group* of size one, was about 39% when applying six regions, while being 13% and 21% for a single region and V4, respectively (similar results appeared in the former section describing the theoretical effects).

Since results may vary depending on the selected regions, performance was also tested over all singleton regions as well as over pairs of regions, triplets etc. (Figure S3). Results varied significantly among singleton regions, yet performance was consistently better and more homogeneous when combining regions *(e.g.,* variability in recall/precision among triplets was smaller than for pairs). Ambiguity also monotonically decreased with increasing the number of regions. However, unlike precision and recall, ambiguity highly depended on the specific selected regions, and displayed more than a two-fold difference between the worst and best combinations.

**Figure 2:**
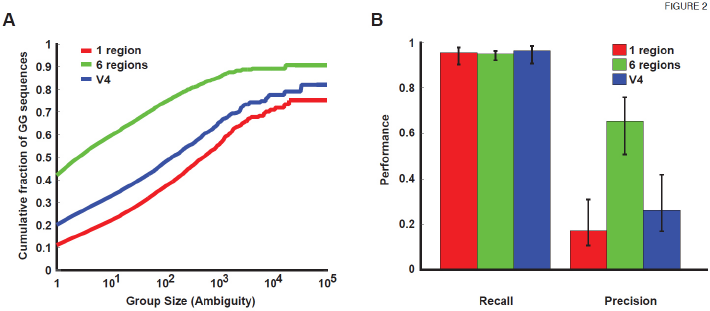
Theoretical resolution and in silico simulation results. (**A**) The *resolution* for one region and six regions. **(B)** Weighted precision/recall of simulated communities in six regions, one region and V4.

## Experimental mock mixture

### Reconstruction accuracy

Figure 3A shows SMURF profiling of our experimental mock mixture, based on one and six regions (results for two to five regions were omitted for clarity and appear in Figure S5). All bacteria were detected, *i.e.,* there were no false negative detections (100% recall), independent of the number of regions applied. There were several false positive detections, mainly *E. faecium* and *P. acnes* which are known contaminations from our lab, and *Wolbachia Pipientis* which was a contamination originating from *Drosophila Melanogaster* samples prepared and sequenced in conjunction. The remaining false positive detections, which may also result from contaminations, sum to about 1%, *i.e.* precision was 99%. For some bacteria frequencies varied between one and six regions (Figure S5). For some bacteria the reconstructed frequency was different from the expected 10% frequency, even after adjusting for the number of 16S rRNA gene copies (Supplementary Results). These differences can be attributed to PCR amplification biases, specifically the lower frequency estimates were related to primers mismatches *(e.g. Eubacterium rectale,* Table S1). Results were almost identical when using the SILVA database instead of the Greengenes database (Figure S6).

### Resolution as function of the number of regions

Figure 3B shows the *ambiguity* of species detected in one and six regions (results for two to five regions appear in Figure S5). *Ambiguity,* measured by the exponent of Shannon's entropy, measures the effective number of 16S rRNA gene sequences that were associated with a specific bacterium. For example the *ambiguity* of *E. coli BL21* was 5384 based on one region and significantly dropped to 25 when using six regions. Hence if we were to follow up on the exact *E. coli* sub-strain we would have had 200 fold less hypotheses to check, as we show later on in the Drosophila Section. Ambiguity was smaller than 100 for all bacteria, and was lower than 10 for six of them, when using the six regions protocol. For comparison, using a single region, *ambiguity* was larger than 100 in nine out of the ten bacteria, while being larger than 1000 for a five of them. Overall, the increase in resolution using six regions was ~100 fold better compared to a single region (Figure 3C).

**Figure 3:**
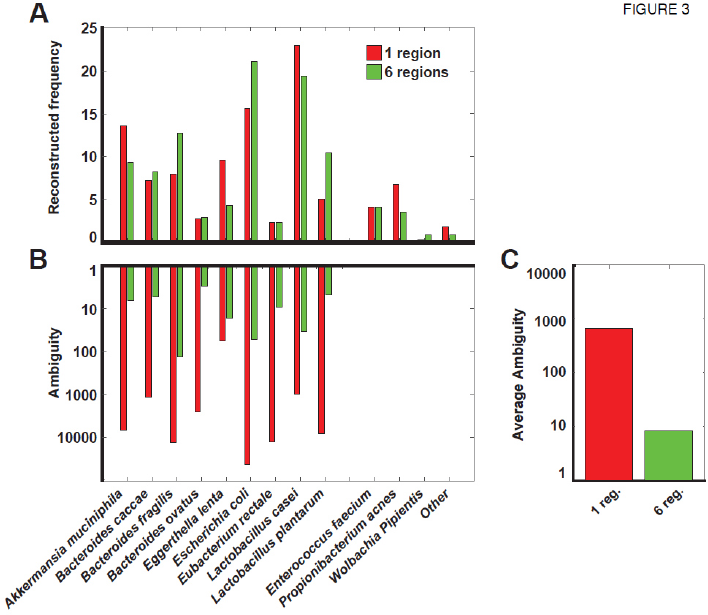
Experimental mock mixture. (**A**) Frequency (percent) of correctly detected bacteria, and of false positive detections (one and six regions). **(B)** Ambiguity as a measure of resolution (one and six regions). **(C)** The average ambiguity (~100 fold smaller for six regions vs. one region).

## Re-analyzing an HMP mock mixture

### Reconstruction accuracy

Figure 4A shows SMURF's profiling results of an HMP mock mixture containing 21 species (SRX 020130). All species were detected in both regions, apart from *Methanobrevibacter* which neither regions amplified [34]. Although mixed so as to create an equal number of 16S rRNA gene copies per bacterium some species were nearly undetectable (e.g. *E. coli and H. pylori)* while others were significantly over represented (e.g. *S. aureus).* As in our mock mixture these biases result either from PCR amplification biases [35] or from specific library preparation steps [21].

### Combining regions increases resolution

As in former cases, the major effect of combining regions was related to increasing resolution. When reconstructing the community jointly from two regions, resolution improved compared to both single regions for eleven bacteria out of twenty detected in the mixture (Figure 4B). For example, *ambiguity* for *H. pylori* using regions V1-V3 and V6-V9, was 137 and 76, respectively. However, combing regions resulted in a significantly smaller *ambiguity,* of seven *H. pylori* 16S rRNA gene sequences, which included the correct sequence. Similarly, for E. *coli* resolution increased from an *ambiguity* of 1892 and 1705 in V1-V3 and V6-V9, respectively, to 602 for the combined solution in which the correct sequence was included. For eight other bacteria resolution of the combined solution improved with respect to one of the regions but was inferior to the other. This is due to sequencing errors and PCR amplification biases, that promote inclusion of sequences with similar amplicon or sequences amplified only in one of the regions to account for the differences in read counts between the regions (see Supplementary Results for an illustrating example). For the bacterium, *A. odontolyticus, ambiguity* was equal (1) for either regions and for their combination, making it invisible on the logarithmic scale. In total, the average *ambiguity* for the V1-V3 and V6-V9 regions was 261 and 216 respectively, while when combining the two regions *ambiguity* reduced to 131 sequences per species, corresponding to a ~2 fold increase in resolution (Figure 4C).

**Figure 4:**
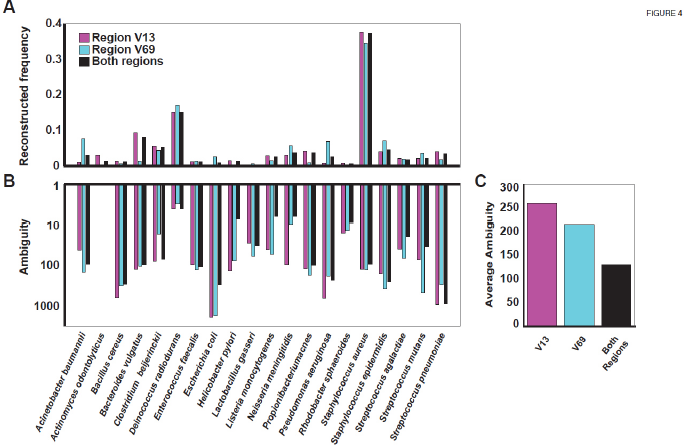
Re-analysis of an HMP 'Even' mock mixture. (**A**) Reconstruction of either V1-V3 or V6-V9 or the two regions jointly. **(B)** Ambiguity when using V1-V3 or V6-V9 regions and by their combination. **(C)** Average ambiguity is reduced ~2 fold when combining the two regions.

## Profiling a Drosophila microbiome

We previously showed that toxic stress in specific tissues of fly larvae can induce developmental phenotypes that persist for multiple generations in non-exposed offspring. An exposure to a G418 toxin led to a delay in larval development, induced the expression of *neoGFP* and modified the adult morphology [36]. The former two phenotypes persisted in subsequent generations of non-exposed offspring.

Since G418 is a broad spectrum antibiotic, this experiment sought to investigate the impact of the G418 toxin on the Drosophila microbial composition and its implications on the inheritance of induced responses (this section presents unpublished data from our project [37]).

## Reconstruction accuracy

Profiling results using one and six regions showed roughly the same microbial composition, namely that G418 lead to a depletion of *Acetobacter* species and to an increase in the frequency of *Lactobacilli* (Figure 5A,B and [37]).

## SMURF enables highly efficient downstream analysis

To check whether the observed change in the host's microbiome played a role in the induction or inheritance of the reported phenotypes, we needed to functionally test whether these phenotypes were induced upon removal of each of the strains present in naïve flies, and whether the inheritance of the phenotypes was interrupted upon reintroduction of these strains to G418 treated flies. Hence, exact strain isolation and identification was required. To achieve that, a set of 16S rRNA gene sequences predicted by SMURF (i.e. hypotheses regarding exact strain identities) were used to design strain-specific PCR/qPCR primers [37], later applied to colonies of individually isolated bacteria in order to identify the various strains. This approach was feasible due to the low *ambiguity* of SMURF's predicted strains, *i.e.,* four, three and 33 for Lactobacillus, Wolbachia and Acetobacter, respectively (Figure 5C, solid). Using a single region there were 887 possible strains for Lactobacillus, 610 for Wolbachia and 130 for Acetobacter (Figure 5C, hollow). Hence, based on a single region such an approach for detecting specific colonies would have been impossible since hundreds of primers should have been used. In such a case one would have to Sanger sequence each of the hundreds of colonies which would have been slow, laborious and quite expensive.

To assess the correctness of SMURF's predictions, we Sanger sequenced the full 16S rRNA gene of bacteria from individual colonies identified by the abovementioned PCR primers to verify their identity. As an example of one such colony, the overlap between the Sanger sequence of a strain of *Lactobacillus* and the predictions based on one and six regions is presented (Fig. 5D, only the top 100 predictions for the single region were shown). One of the sequences indeed correctly identified the strain, using both one and six regions, while the rest were wrong. Figure 5D emphasizes a tremendous advantage of the multiple regions approach whose number of false positive detections was orders of magnitude smaller, thus allowing identification of the specific strain present in the sample quickly and efficiently.

**Figure 5.**
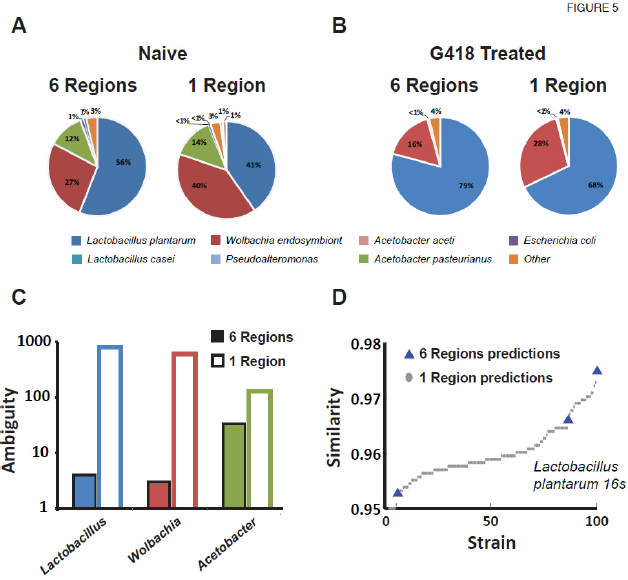
Reconstruction of bacterial populations in *D. melanogaster* following toxic treatment. **(A)** Naïve flies using six regions (left) and one region (right). **(B)** Same for flies reared on medium containing the G418 toxin. **(C)** Ambiguity of the three most abundant species. Profiling based on a single region (hollow bars) and six regions (full bars). **(D)** Similarity between L. *plantarum* predicted strains and the Sanger sequence of the strain isolated from the detected colony (top 100 sequences).

Finally we used the different isolated bacteria to functionally test the strains and to uncover two strains having an interesting biological activity [37]. The G418 toxin led to a heritable selective depletion of commensal *Acetobacter*s and explained the heritable developmental delay. Reintroducing *Acetobacter*s, detected by SMURF, prevented this inheritance.

## Discussion

The two main approaches for inferring the composition of an unknown mixture of bacteria are either 16S rRNA gene sequencing or whole genome shotgun sequencing. Although sequencing whole genomes provides much more information and does not require bias-prone PCR amplification, sequencing the 16S rRNA gene has several advantages. First, most computational methods for analyzing sequencing results of either the 16S rRNA gene or of whole genome sequencing rely on a database of sequences. Although current databases of bacterial genomes are growing fast, they still contain about an order of magnitude less sequences than 16S rRNA gene databases. Consequently, a species may be identified when sequencing the 16S rRNA gene while being missed when analyzing shotgun sequencing results since its genome has not yet been deposited in the databases. A second potential advantage of 16S rRNA gene based methods is related to profiling samples having a low bacterial biomass which makes it imperative to amplify bacteria-related sequences, and renders shotgun methods highly inefficient *(e.g.,* sequencing without prior amplification would result in an overwhelming number reads originating from the host's DNA and almost no bacterial sequences). Third, 16S rRNA gene sequencing allows for multiplexing of far more samples per lane compared to shotgun sequencing, and thus yields a much lower cost per sample. Hence, we view 16S rRNA gene based profiling as a complementary approach to whole genome sequencing, where each approach should be chosen according to the experimental setting.

In this work we focused on 16S rRNA gene sequencing and sought to improve upon the state of the art in the field. Sequencing the 16S rRNA gene faces three main challenges, that eventually hamper the accuracy of profiling and its reproducibility. The first challenge is limited resolution of current primer pairs, *i.e.,* many bacteria share the same sequence over the amplicon and thus the exact identity of bacteria in the community cannot be inferred. The second challenge is 'partial' primer universality, *i.e.,* different primers pairs amplify certain subsets of all bacterial 16S rRNA gene sequences [18–20] and hence primer choice governs profiling results. Primers may also be biased towards amplifying certain genera because of the sequence composition of the database used for their design [38]. The third challenge in 16S rRNA gene profiling is PCR-related errors and biases, which may result from several sources, e.g., primers' mismatches [35], adding sample indices (i.e., 'barcodes') to the primer sequence [21, 39], or other library preparation steps [21].

In this work we present a novel analysis framework termed SMURF that directly addresses the first two problems via computationally combining multiple 16S rRNA gene short amplicon sequencing data, and thus improving upon current best approaches. The last issue of sample preparation biases is only indirectly addressed in this work. Although combining results from different regions tends to average the effects of such biases and errors, it is certainly beneficial to carefully choose the set of primers (Supplementary Figure S3) and to modify library preparation procedures so as to reduce such errors and biases [21].

The set of six primer pairs used in this study should be considered a first trial that served for presenting the advantages of combining regions. Since accurate profiling evidently depends on the properties of the chosen set of primers, it would be interesting to seek an optimal set. Also, it should be noted that accuracy and resolution are limited by the number of sequencing reads. Hence, in order to assure accurate profiling of low frequency bacteria a sufficient number of reads should be allocated to each region. The latter also dictates the number of samples that may be multiplexed per lane.

## Conclusions

Short Multiple Regions Framework (SMURF) represents a new approach in bacterial community reconstruction. A significant improvement in resolution is achieved as has been displayed theoretically, using *in silico* simulations and by analyzing two mock mixtures and real-life samples.

In many cases microbial profiling at the genus or even species level may be achieved using standard primers (e.g. V4 primers) and does not require combining regions. However, in cases where downstream analyses are required, increasing resolution via SMURF, i.e., reducing the reported number of full-length 16S rRNA gene sequences per profiled bacterium is crucial as demonstrated by our *D. melanogaster* experiments.

SMURF can be applied to standard sample preparation with very little modifications at the PCR stage *(i.e.,* sample preparation is independently performed using each primer pair), and allow re-analysis of samples that were profiled using several primer pairs. Apart from increased resolution, effects of PCR biases will tend to be 'averaged' and hence be less pronounced. Also, the basic algorithm which is an adaptation of [12], allows fast convergence and avoids the necessity to align reads, and hence the computational overhead in applying SMURF is small.

SMURF paves the way for high resolution profiling of fragmented DNA in low biomass samples, *e.g.,* FFPE blocks, which are abundant and relatively easy to acquire (compared to snap frozen samples); hence SMURF may be highly beneficial in, e.g., cancer microbiome studies. Additionally, it would be beneficial to optimize a set of primer pairs so as to support a multiplexed PCR reaction. Although such optimization requires significant one-time labor, it would eventually save precious material and allow highly efficient profiling.

SMURF is not restricted to amplicons of the 16S rRNA gene and can be seamlessly adapted to combine any set of amplicons, e.g., in Multilocus Sequence Typing (MLST). Such profiling may be required when the 16S rRNA gene is not sufficient to provide the required resolution.

## Declarations

### Ethics approval and consent to participate

Not applicable.

### Consent for publication

Not applicable.

### Availability of data and material

Experimental data are available at the MG-RAST server

(http://metagenomics.anl.gov/linkin.cgi?project=mgp20457)

## Authors' contributions

AZ, AA, GF and NS conceived and planned the study. AZ and ME designed and performed the experiments. GF and NS devised the algorithm and analyzed the data. GF and NS drafted the manuscript with input from YS and PJT, and all authors critically revised and approved the final version of the manuscript.

## Acknowledgements

The authors would like to thank Henry J. Haiser for preparing the bacterial mock mixture.

## Additional materials

Supplementary Results.docx

## Supplementary Results

### Theoretical effect of combining short regions

The *group* size of each bacterial sequence in the Greengenes (GG) database, namely the number of GG sequences that share the same sequence over the relevant amplified regions, was calculated. Figure S1 shows the fraction of GG sequences that belong to a *group* of up to a certain size, for one to six regions and for the V4 region. Using two regions was sufficient for better performance than V4. The sixth region seemed to provide only a small advantage over using the first five regions.

**Figure S1.**
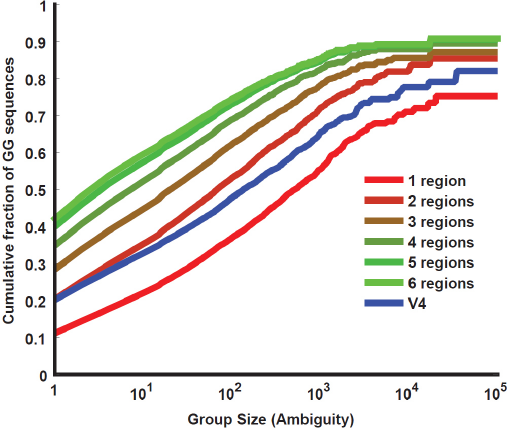
(related to Figure 2A): Theoretical resolution. Each sequence in the Greengenes (GG) database was assigned to a *'group', i.e.,* the set of GG sequences from which it is indistinguishable over the relevant amplified region(s). The figure shows the fraction of GG sequences that belong to a *group* of up to a certain size for one to six regions and for V4. The *group's* size coincides with the *ambiguity* in this case and hence measures the resolution by which a bacterium may be identified.

### *In silico* simulations

A thousand mixtures were selected, each comprising one hundred bacteria randomly selected from the GG database and assigned a power law frequency distribution. Weighted precision and weighted recall were plotted for one to six regions and for using V4 primers, as a function of the *total* number of reads applied. The average number of reads per region was the total number of reads divided by the number of regions. Weighted precision monotonically improved when increasing the number of regions (two regions were sufficient to provide comparable precision to that of V4). Weighted recall was high in all cases, when the total number of reads was high enough *(e.g.,* higher than 200,000). However, when allocating a low number of reads weighted recall decreased. This reduction in performance was more significant for higher number of regions, since the number of reads per region was insufficient to detect the low frequency bacteria.

**Figure S2.**
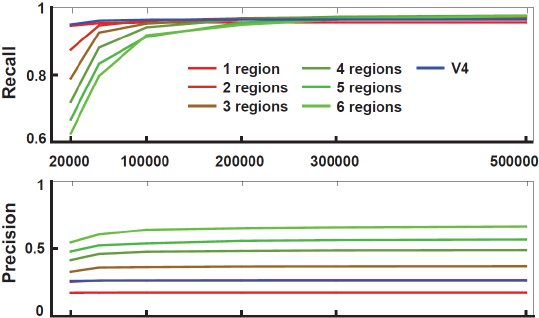
(related to Figure 2B): Simulation results. Performance, *i.e.,* weighted precision and weighted recall of simulated communities for one to six regions and for V4 as function of total number of reads. Error bars were omitted for clarity.

### Combinations of regions and their effect on mixture reconstruction

This section presents results across all possible subsets of regions up to six (as opposed to the former section that considered a single set of regions for each size). For each set of regions, 1000 mixtures were selected similarly to former simulations (using a total 200,000 reads across all regions). Figure S3 shows weighted recall, weighted precision and ambiguity across all sets of regions, segregated into singletons, pairs of regions, triplets, *etc̤* The 'size' of a set of regions refers to the number of combined regions.

*Weighted recall and precision:* Singleton regions displayed highly variable recall values, corresponding to the difference in universality among them. Weighted recall and precision for singleton regions were inferior to those of sets of regions. Weighted recall for all larger sets was high, while slightly decreasing for larger sets due to a lower number of reads per region. Weighted precision monotonically improved when increasing the number of regions. The variability in recall and precision among all combinations of regions of a certain size (e.g., among pairs) decreased with increasing the set size.

*Ambiguity:* Ambiguity was highly variable across different sets of regions, significantly decreasing when increasing the number of regions. Interestingly, ambiguity among sets of regions of the same size may vary by more than two fold. Although this phenomenon reflects the properties of the specific set of primer pairs, it shows that profiling would certainly benefit from a careful selection of sets of primers. In addition, in almost all cases it was favorable to use all possible primers rather than any subset of them.

**Figure S3.**
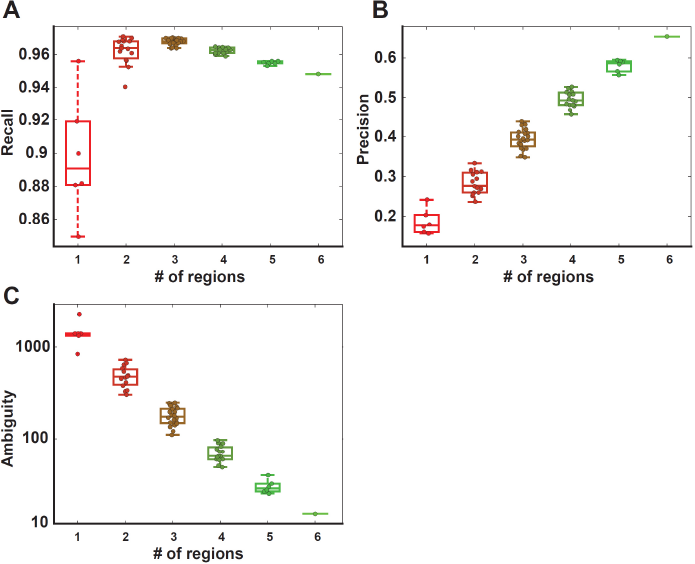
Results across all subsets of regions. Performance, *i.e.,* weighted recall (panel A), weighted precision (panel B) and ambiguity (panel C) of simulated communities for sets of one to six regions. Ambiguity is shown on a logarithmic scale.

### Read error effect

Performance was tested for a range of assumed constant *p_e_* values. For low values of *p_e_*, the algorithm implicitly assumes that all reads are correct, and hence more bacteria appear in the reconstructed mixture and precision deteriorates. For higher values of p_e_, the algorithm was able to correctly identify some of the noisy reads as such, which results in less falsely detected bacteria. Performance was robust for a wide range of p_e_ values (Figure S4). Even under an extreme assumption of p_e_ = 0.001 the degradation in precision was less than 3%. Based on these observations (and (Ross et al. 2013)) we chose to use *p_e_* = 0.005 for reconstruction of both simulated and experimental mixtures.

**Figure S4.**
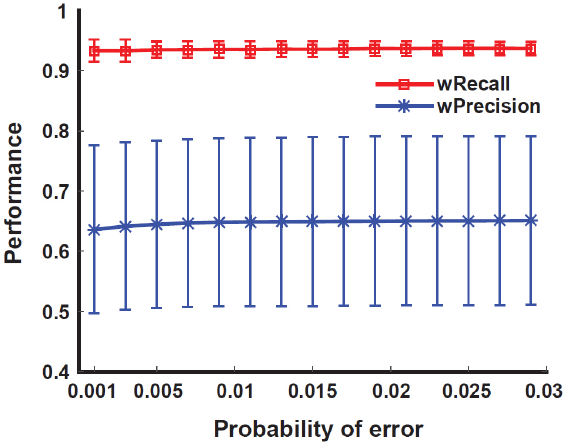
Weighted precision and recall averaged over 100 simulated communities as function of the value assumed for the sequencing error rate. *p_e_*.

## Experimental mock mixture - normalizing the number of 16S rRNA gene opérons

Table S1 compares reconstruction results based on six regions with and without adjusting for the number of 16S rRNA gene operons (operons for each bacterium were extracted by mapping the primers on its whole genome sequence). To compare results before and after normalization, the contaminations and false positive detections were excluded, and frequencies were renormalized to 1 in both cases. Operon adjusted reconstruction was closer to the original uniform distribution (Shannon diversity increased from 7.57 sequences to 8.94 sequences following operon adjustment), although for some bacteria the estimated frequency significantly deviated from the original proportion of 10%. This may be attributed to variable PCR amplification efficiency due to primer mismatches (the mixture was assembled using pre-extracted quantified DNA and hence extraction efficiency did not contribute to variable frequencies).

**Table S1:** Experimental mock mixture reconstruction results before and after operon correction. For each bacterium the number of operons identified in their whole genome sequence is shown together with the reconstructed frequency before and after adjustment for the number of operons, based on six regions.

| Mock mixture bacterial species | # of operons | Not adjusted | Adjusted |
| --- | --- | --- | --- |
| *Escherichia coli* BL21 | 7 | 23.29 | 14.01 |
| *Lactobacillus casei* | 5 | 21.38 | 18.0 |
| *Bacteroides fragilis* | 6 | 14.02 | 9.84 |
| *Lactobacillus plantarum* | 5 | 11.49 | 9.67 |
| *Akkermansia muciniphila* | 3 | 10.26 | 14.4 |
| *Bacteroides caccae* | 5 | 9.06 | 7.63 |
| *Bacteroides ovatus* | 3 | 3.24 | 4.55 |
| *Eubacterium rectale* | 5 | 2.56 | 2.16 |
| *Eggerthella lenta* FAA 1-3-56 | 1 | 2.35 | 9.87 |
| *Eggerthella lenta* DSM2243 | 1 | 2.35 | 9.87 |

## Experimental mock mixture

### Reconstruction accuracy

SMURF profiling of our experimental mock mixture was performed based on one to six regions. For most bacteria frequency seemed to 'stabilize' for two or more regions (Figure S5A). Also, the fraction of 'other' false positive detections decreased from ~2% to ~1% when using three or more region.

### Resolution as function of the number of regions

*Ambiguity* of species detected in one to six regions was measured by the exponent of Shannon's entropy (Figure S5B). Although all bacterial species were correctly detected for any number of regions and bacterial frequency seemed to stabilize even for two regions, in most cases resolution continued to improve as the number of profiled regions was increased. Due to inherent noise in the reconstruction process the resolution of a specific bacterium may not be strictly monotonic with increasing the number of regions. For example, the average group size of *B. ovatus* increased due to noise when we added the fourth region, but it improved again significantly when adding of the fifth region. An artificial example of such an effect is given below.

**Figure S5.**
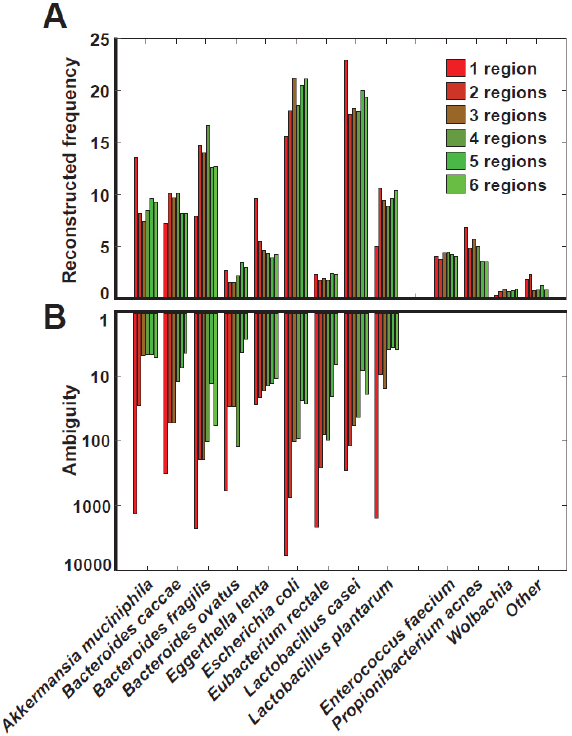
(related to Figure 3) Experimental mock mixture. (A) Reconstruction results: Frequency (percent) of the reconstructed bacteria, both correct and false positive detections. (B) *Ambiguity* as a proxy for resolution: the exponent of Shannon's entropy of each bacterium in the mock mixture is shown, for one to six regions on a logarithmic scale.

### SMURF's results are robust to the selected 16S rRNA gene database

SMURF's reconstruction performed using the SILVA database^1^ (Quast et al. 2013) was highly similar to the results obtained for the Greengenes-based SMURF (Figure S6). Results display robustness to the specific choice of sequences' database.

**Figure S6.**
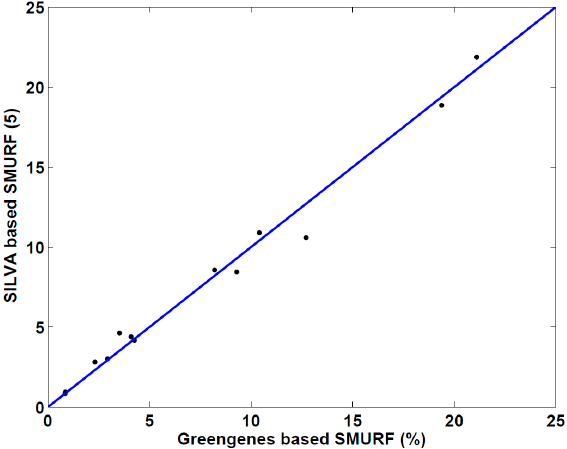
Database dependence: A scatter plot of SMURF's mock mixture reconstruction (in percent) based on six regions: for a Greengenes-based SMURF (horizontal) vs. a SILVA-based SMURF (vertical). Each dot corresponds to a mock mixture bacterium.

### The effect of noise on resolution

The following synthetic example describes a scenario in which noise *(i.e.,* PCR amplification errors and sequencing errors) causes a reduction in resolution when adding a region.

*The ad hoc database:* The 'experiment' is performed using two regions “1” and “2”. The database includes a bacterium *A* which is amplified by both primers' of region “1” and of region “2”, and ten additional bacteria, *B, C,…, K,L,* which are amplified only in region “2”. The latter ten bacteria have the same sequence as bacterium *A* in region “2”.

Hence the matrix *M* for a single region is:

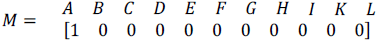

And for two regions, the matrix is:

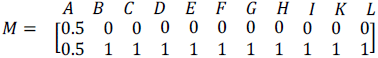

In this synthetic example we assume that *p_e_* = 0 and thus *Q* = *M*.

*The 'mixture':* Given the matrix *M* we examine the results of profiling a 'mixture' containing a single bacterium *A*. Profiling is performed using region “1” and using both region “1” and region “2”. We first consider the noiseless case and then introduce noise.

*'Experimental' measurement - the noiseless case:* In the absence of noise and since the mixture includes only bacterium *A*, the measurement vector *y* for profiling based on only region “1” is given by *y* = 1. The vector *y* for profiling based on two regions is given by: 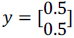.

The solution of the optimization problem for a single region and for two regions is the same in this case, and includes only bacterium *A*:

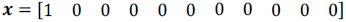

Resolution *(i.e. ambiguity)* in both cases is 1.

*Experimental' measurement - the noisy case:* When profiling is based on a single region, noise has no effect since y = 1.

Hence, ***x***=[1 000000000 0] and *ambiguity* is 1, as in the noiseless case.

However, assume that due to noise, the measurement vector for two regions slightly deviates from its noiseless value:

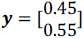

The solution of the optimization problem in this case would be

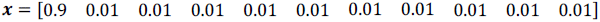

And hence the *ambiguity* is 1.74, which is larger than for a single region.

## Extended legends for manuscript’s figures

**Figure 1: A Schematic description of SMURF. (A)** A schematic comparison between single and multiple region profilings. The two bacterial species (mocca and light gray) in the mixture are indistinguishable when profiling is based on a single (red) region. Moreover the existance of a third (light blue) bacterial species may not be ruled out. In contrast, when profiling is based on multiple regions (two in this case) the true composion may be reconstructed. **(B)** The SMURF flow diagram describing the internal steps of a typical analysis.

**Figure 2: Theoretical resolution and in silico simulation results. (A)** Theoretical resolution. The fraction of GG sequences that belong to a ‘group’ (*i.e.*, the set of GG sequences that are indistinguishable over the relevant region) of up to a certain size for three cases: our single region, V4 and our six regions. Using six regions provided a significant increase in resolution, namely more sequences belonged to smaller groups, and hence may be better identified. The group’s size coincides with the ambiguity in this case. **(B)** In silico simulation results. Weighted precision and weighted recall of simulated communities in three cases: six regions, a single region and V4. Error bars represent up/down mean absolute deviation over 1000 simulated communities.

**Figure 3: Experimental mock mixture. (A)** Reconstruction results: Frequency (percent) of correctly detected bacteria, and of false positive detections. Profiling was performed using a single region and using all six regions (legend applies to all panels). **(B)** Ambiguity as a measure of resolution: The exponent of Shannon’s entropy of each bacterium in the mock mixture, shown for one and for six regions on a logarithmic scale. **(C)** The average ambiguity, which is ~100 fold smaller when using six regions compared to a single region.

**Figure 4: Re-analysis of HMP data. (A)** Reconstruction of the HMP ‘Even’ mock mixture (SRX 020130) from either V1-V3 or V6-V9 regions and from the two regions jointly (legend applies to all panels). **(B)** Ambiguity of profiling the HMP mock mixture: The exponent of Shannon’s entropy of each bacterium in the HMP mock mixture SRX 020130, profiled either using V1-V3 or V6-V9 regions and by their combination. **(C)** Average ambiguity is reduced ~2 fold when combining the two regions.

**Figure 5. Reconstruction of bacterial populations in D. melanogaster following toxic treatment. (A)** Reconstruction of the bacterial populations at the species level in naïve flies using six regions (left) and one region (right). **(B)** Same for flies reared on medium containing the G418 toxin. Overall profiling at the species level was highly similar between one and six regions. **(C)** Ambiguity of the three most abundant species. Profiling based on a single region (hollow bars) resulted in much higher species ambiguity than for six regions (full bars), which allowed for much more efficient downstream analysis. **(D)** Similarity between *L. plantarum* predicted strains and the Sanger sequence of the strain isolated from the detected colony (the top 100 sequences are shown). The correct strain appeared in the predicted set for both one and six regions (dots and triangles, respectively), although the number of other L. plantarum false predictions were much larger in the case of a single region.

## Footnotes

1 The SILVA version 128 resulted in a database of 2,014,374 16S rRNA gene sequences after accounting for sequence nucleotide ambiguities, and discarding sequences whose length was either longer than 2500bp or shorter than 1200bp.

